# Extrafollicular plasma cells disable dendritic cell-T-cell priming in tumor-draining lymph nodes

**DOI:** 10.64898/2026.04.20.719651

**Authors:** Elena Alberts, Victoire Boulat, Miu Shing Hung, An Qi Xu, Jelmar Quist, Mengyuan Li, Fangfang Liu, Isobelle Wall, Gregory Verghese, Carin Andrea Brundin, Ananya Bhalla, Mats Jönsson, Carlos Castellanos, James Rosekilly, Cheryl Gillett, Johan Staaf, Caetano Reis e Sousa, Sophia N. Karagiannis, Anita Grigoriadis, Dinis Pedro Calado

## Abstract

How plasma cells (PCs) shape anti-tumor immunity is unclear. We hypothesized that conflicting prognostic associations reflect differences in immune context and PC ontogeny. We identify extrafollicular (EF)-PCs as an antibody-independent checkpoint that aborts priming by disabling the cDC1→CD8^+^ T-cell axis in tumor-draining lymph nodes (td-LNs). EF-PCs blunt cDC1 activation and CCR7-guided repositioning into T-cell zones, precluding formation of TCF1⁺ stem-like CD8⁺ T-cells. Depleting EF-PCs in vivo restores cDC1 trafficking, expands the stem-like reservoir, increases intratumoral CD8⁺ infiltration, and restrains tumor growth; benefit is lost with CD8 T-cell ablation. Neither serum transfer nor Fcγ receptor blockade reverses tumor control, supporting a non-canonical, antibody-independent mechanism. Across independent triple-negative breast cancer cohorts, we find EF-PC hyperplasia in td-LNs and tumors; and within immune-cold cases, EF-PC burden stratifies poor prognosis and metastatic risk. A cross-species EF-PC signature maps to a conserved PC-state across cancer types that is linked to poor outcome and immune-checkpoint blockade resistance. EF-PCs thus relocate the dominant failure point to td-LNs and offer a tractable upstream target to convert immune-cold tumors into immune-responsive disease.

## Main

Plasma cells (PCs) are classically defined as antibody-secreting factories, so their influence in cancers has largely been inferred from serology and FcγR-biology. In contrast, antibody-independent functions are less investigated, and pathological studies enumerating PCs in the tumor bed have reported heterogenous associations with outcome^1–10^. These observations suggest that greater granularity is needed to define when, where, and which PCs matter.

Two axes likely account for much of the observed heterogeneity. First, immune context; many cohorts are not stratified by tumor-infiltrating lymphocyte (TIL) density, conflating immune-hot with immune-cold disease in which PC functions may be context-dependent or even opposing. Triple-negative breast cancer (TNBC), the most aggressive and therapeutically refractory breast cancer, exemplifies the importance of tumor context^11^. TILs are a strong, guideline-recognized prognostic biomarker, and tertiary lymphoid structure (TLS) and B-cell-rich microenvironments correlate with immune checkpoint blockade (ICB) response^12,13^. However, the majority (>80%) of TNBCs are immune-cold, displaying poor response to ICB and worse outcomes^14,15^.

Second, ontogeny. PCs arise via germinal-center (GC) or extrafollicular (EF) programs that differ in kinetics, anatomical localization, and selection history. In GCs, follicular B-cells undergo iterative somatic hypermutation (SHM) with stringent selection, producing affinity-matured PCs. EF responses, typically centered in lymph node (LN) medullary cords, are rapid and proliferative, with limited SHM and selection, and therefore produce PCs with minimal affinity maturation. EF responses are a normal early arm of humoral defense but can become maladaptive in chronic settings as reported in persistent infection and autoimmunity^16,17^. These origin-specific features plausibly confer distinct, and at times opposing effects on anti-tumor immunity. Recent cross-cancer atlases have mapped intratumoral PC states; however, their functional roles remain to be defined^18,19^. Addressing this gap requires looking beyond the tumor bed to secondary lymphoid organs, the principal sites of PC generation, especially tumor-draining lymph nodes (tdLNs).

Here we resolve both issues. Across independent TNBC cohorts; Guy’s (n=124)^20^, SCAN-B (n=202)^21,22^ and METABRIC (n=101)^23^; we stratified tumors by immune context using TIL-defined immune-hot versus immune-cold and determined PC ontogeny. This revealed cancer-induced hyperplasia of EF-PCs (low SHM/high proliferation) at the tumor bed and within medullary cords of matched td-LNs. EF-PC burden defined a biomarker that identified the poor-prognosis subset within immune-cold tumors and tracked metastatic risk.

To isolate EF-PC function without pan-B-cell confounding effects, we used three orthogonal PC-targeted strategies (αCD138, Jchain_CreERT2-DTA^24,25^, and bortezomib). Mechanistically, EF-PCs co-localized with DCs in td-LNs and imposed an antibody-independent, non-canonical checkpoint that dampened cDC1 activation and CCR7-dependent relocation to T-cell zones, aborting CD8⁺ T-cell priming. EF-PC depletion restored DC→T crosstalk, expanded TCF1⁺ stem-like memory CD8⁺ T-cells, boosted intratumoral effectors, and restrained tumor growth; functionally converting immune-cold into immune-hot tumors. A cross-species EF-PC transcriptional signature mapped to poor-prognosis, immunotherapy-resistant PC states across TNBC, HER2⁺ breast, basal cell, clear-cell renal, colorectal and lung cancers, nominating EF-PCs as druggable checkpoints upstream of the tumor. Collectively, these data revealed an EF-PC→DC→T-cell circuit in cancer immunity, highlighting a tractable avenue to reinvigorate anti-tumor immunity in TNBC and other immune-excluded cancers.

### Intratumor PC burden identifies high-risk immune-cold TNBC

We profiled at the tumor bed intratumoral B-cell populations across independent TNBC cohorts, Guy’s (n=124)^20^, SCAN-B (n=202)^21,22^, METABRIC (n=101)^23^ by immune deconvolution of bulk tumor transcriptomes with a scRNA-seq-derived reference^26^ (**Fig. 1A**). In accord with the International TILs Working Group criteria^27^ and using histological scoring (Guy’s and SCAN-B) and computational estimation (METABRIC), tumors were stratified by stromal TIL density (sTIL ≤10% vs >10%) to separate immune-cold from immune-hot disease, respectively. Analysis of the Guy’s cohort using multivariate analyses adjusted for age and clinical covariates (tumor size, grade, stage, nodal involvement and treatment status) showed that memory and GC B-cell signatures associated with reduced distant metastasis risk, both in the unstratified set and in immune-hot (high-sTIL) tumors, in agreement with previous work^28^ (**Fig. 1B**). By contrast, the PC signature was not prognostic in the unstratified or immune-hot (high-sTIL) tumors, but within immune-cold (low-sTIL) tumors it associated with increased hazard of distant metastasis (**Fig. 1B**).

**Figure 1:**
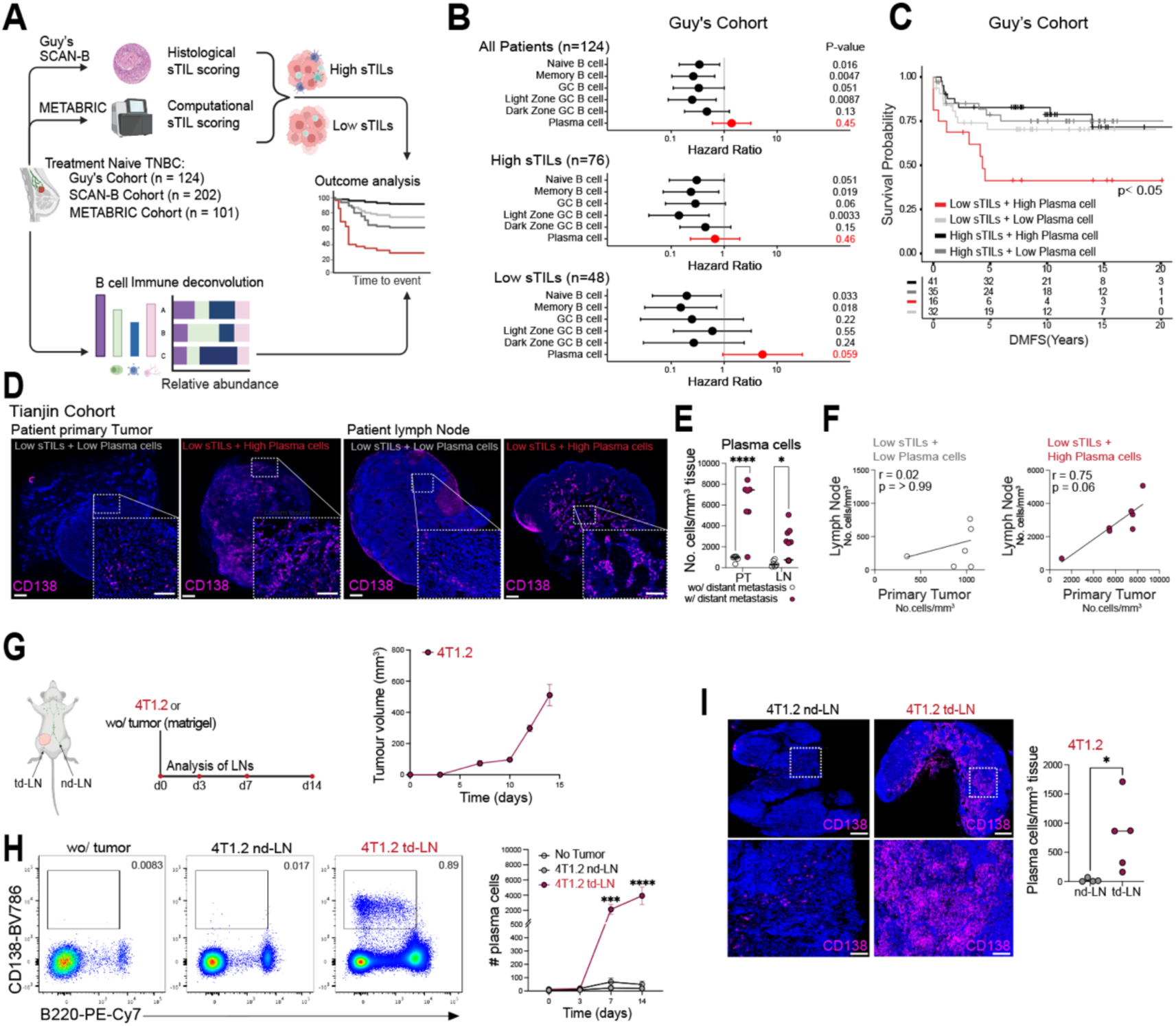
PC enrichment identifies poor-prognosis within “immune-cold” TNBC patients. **(A)** Schematic overview of patient TNBC cohorts, stromal tumor-infiltrating lymphocyte (sTIL) scoring methods, and B-cell-focused immune deconvolution used for outcome analysis. **(B)** Multivariate analysis of the Guy’s TNBC cohort (n = 124), assessing key B-cell populations across all patients and stratified by sTIL scores. Analyses were adjusted for age, tumor size, tumor stage, and nodal stage. **(C)** Kaplan–Meier analysis of distant metastasis-free survival (DMFS) in the Guy’s cohort. Patients are stratified by sTIL level and plasma cell gene signature expression: high sTIL/high signature (black), high sTIL/low signature (dark grey), low sTIL/low signature (light grey), and low sTIL/high signature (red). **(D)** Representative immunofluorescence images of CD138⁺ plasma cells in primary tumors and involved lymph nodes from two low sTIL patients in the Tianjin cohort. Scale bar: 1mM. **(E)** Quantification of CD138⁺ plasma cells in primary tumors (PT) and involved lymph nodes (in-LN) comparing patients with or without distant metastasis. Each data point represents a single PT or in-LN. **(F)** Correlation between CD138⁺ plasma cell infiltration in in-LNs and matched PTs of patients without (left) or with (right) distant metastasis. **(G)** Schematic of TNBC mouse models used, with timeline of tumor and lymph node harvesting post-tumor inoculation, and tumor growth curve. **(H)** Flow cytometry plots and quantification of CD138⁺B220^neg^ plasma cells in LNs from tumor-free control mice (No Tumor), non-draining (nd-LN) and tumor-draining lymph nodes (td-LN) from mice bearing 4T1.2 tumors over time, shown as absolute numbers. **(I)** Immunofluorescence images and quantification of CD138⁺ plasma cells in nd-LNs and td-LNs from the 4T1.2 mouse model, expressed as plasma cells per mm² of tissue. Scale bar top panel: 500μm; scale bar bottom panel: 100μm. **Statistical analysis:** Survival curves (C) were compared using the Likelihood Ratio test; HR = hazard ratio, CI = confidence interval. Comparisons in (E) and (I) used the Mann–Whitney U test *p* < 0.05 (\**),p < 0.001 (*\*\*\**), p < 0.0001 (***\*\*\*\***). Spearman’s rank correlation was used for (F). Data in (H) are data representative of five independent experiments.

Kaplan-Meier analyses mirrored these findings, within immune-cold (low-sTIL) tumors, high PC signature tracked with significantly shorter distant metastasis free survival (DMFS) in the Guy’s cohort and reproduced in both METABRIC and SCAN-B cohorts (**Fig. 1C**; **Ext. Fig. 1A**). Notably, immune-cold (low-sTIL) tumors with low PC signal achieved DMFS comparable to immune-hot (high-sTIL) tumors (**Fig. 1C**), indicating that the absence of PC activity mitigates the otherwise adverse prognosis of immune-cold disease.

To validate the transcriptomic signal of PC infiltration, we performed CD138 staining on 37 primary tumors from the SCAN-B cohort, selecting cases transcriptionally classified as immune-cold (low sTIL) with either low or high PC signature. Pathology confirmed higher PC density in signature-high tumors (**Ext. Fig. 1B,C**), supporting the deconvolution results and indicating that PC enrichment within an immune-cold microenvironment marks a clinically high-risk subset of TNBC.

### Concordant PC burden between tumor and matched LN

We next asked whether PC infiltration at the tumor bed reflects a broader immunopathological program involving LNs. In an independent TNBC cohort (Tianjin) with 16 primary tumors matched to involved LNs, CD138⁺BLIMP1⁺ PCs were present in both compartments (**Fig. 1D**; **Ext. Fig. 1D**). High PC infiltration in the primary tumor and in the paired LN associated with distant metastasis (**Fig. 1E**). Notably, PC densities in tumor and matched LN were correlated only in patients who developed metastasis (**Fig. 1F**), consistent with a coordinated expansion across sites and suggesting that td-LNs may serve as active sites of PC differentiation.

To test whether immune-cold TNBCs drive PC hyperplasia in td-LNs, we studied an immune-cold ICB-refractory TNBC model, 4T1.2^29–31^. These cancer cells are derived from an aggressive lung-metastatic clone of 4T1. Following orthotopic implantation, we collected td-LNs and contralateral non-draining LNs (nd-LNs) at serial time points (**Fig. 1G**). As expected, 4T1.2 tumors expanded rapidly from day 10 post-inoculation (**Fig. 1G**) and td-LNs were consistently larger than nd-LNs and LNs from tumor-free mice (**Fig. 1H**; **Ext. Fig. 1E**). Pronounced PC hyperplasia emerged in td-LNs as early as day 7, mirroring high-risk immune-cold human TNBCs (**Fig. 1H**). Immunofluorescence localized these PCs predominantly to medullary cords (**Fig. 1I**). Together, these data indicate that td-LNs undergo tumor-driven remodeling that promotes PC hyperplasia, a program linked to poor-prognosis in immune-cold TNBC.

### Cancer driven EF-PC hyperplasia within LNs

4T1.2 tumors also elicited GC B-cell responses in td-LNs, however, the GC reaction peaked by day 7 and declined rapidly thereafter **Ext. Fig. 2A,B**, whereas PCs continued to expand and remained elevated at day 14 (**Fig. 1H**). Across td-LNs, PC abundance inversely correlated with GC B-cell frequency (**Ext. Fig. 2C**), indicating tumor-driven skewing toward extrafollicular (EF)-PC generation, uncoupled from GC selection. Consistently, in human immune-cold TNBC, lymph nodes from patients with immune-cold tumors with high PC burden (PC^High^) contained fewer GC structures and showed a redistribution of PCs to EF zones compared with immune-cold tumors with low PC burden (PC^Low^) (**Fig. 2A,B**), supporting an EF origin of the PC response (**Fig. 2C**).

**Figure 2:**
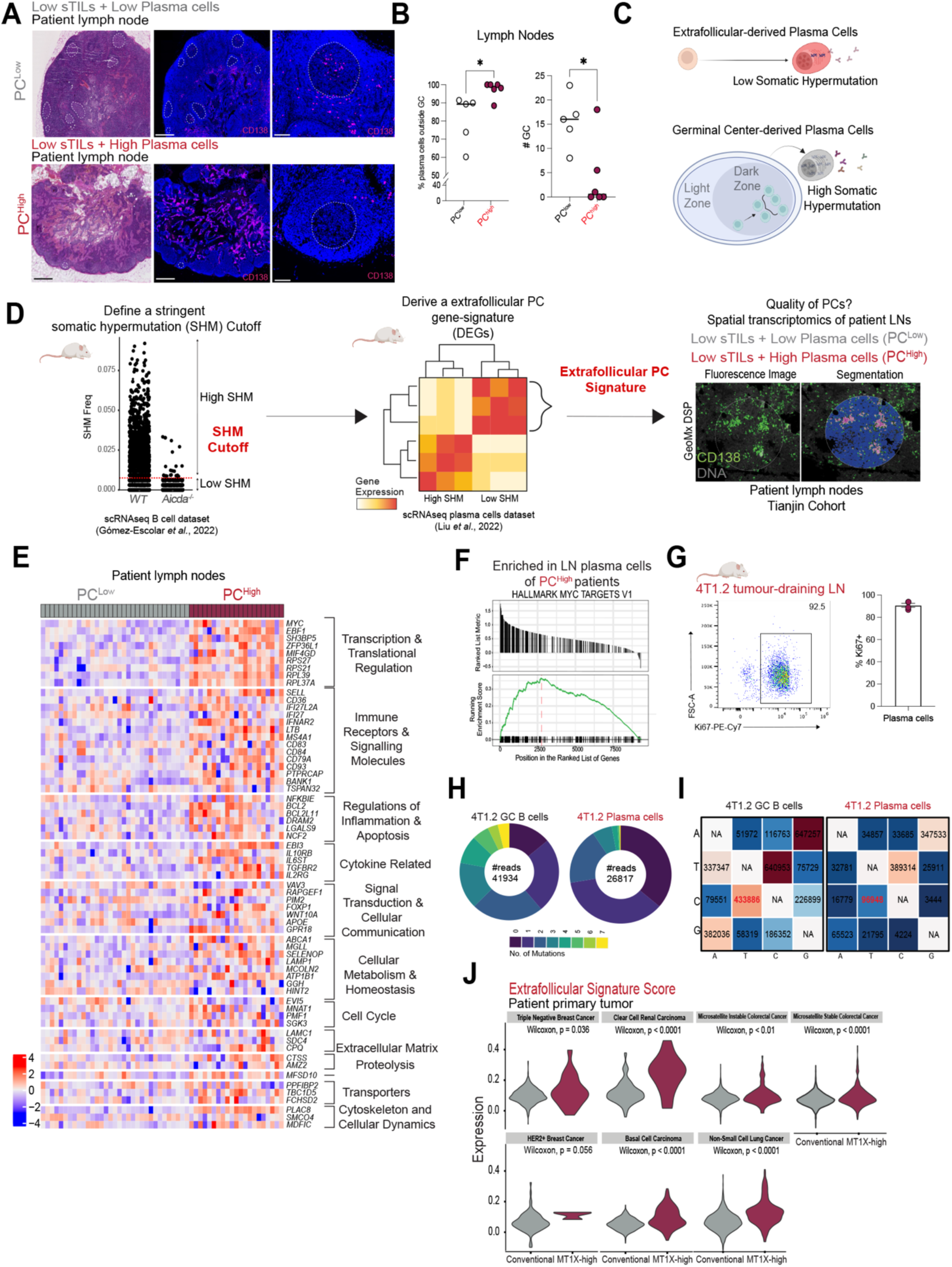
Cancer driven extrafollicular-derived PC hyperplasia in LNs. **(A)** Representative H&E and CD138 immunofluorescence images of lymph nodes from low sTIL TNBC patients with either high (PC^high^) or low (PC^low^) plasma cells. Dashed white lines indicate germinal centres. Scale bar left/middle panel: 800μm, scale bar right panel: 100μm. **(B)** Quantification of the proportion of CD138⁺ plasma cells located outside germinal centers (GCs), and total GC numbers, in PC^low^ vs PC^high^ patients. **(C)** Schematic overview of plasma cell differentiation pathways, highlighting extrafollicular (EF) and GC-derived routes. **(D)** Strategy for defining an EF-plasma cell gene signature: public mouse single cell datasets were used to identify EF-specific genes, which were then applied to spatial transcriptomic (Nanostring GeoMx DSP) data from human cancer-associated lymph node plasma cells. **(E)** Heatmap showing EF signature expression in plasma cells from lymph nodes of PC^low^ and PC^high^ patients. **(F)** Gene Set Enrichment Analysis (GSEA) for the Hallmark “Myc Targets V1” pathway in plasma cells from PC^high^ patients. **(G)** Representative flow cytometry of Ki67+ plasma cells from td-LNs of 4T1.2 tumor-bearing WT mice 14 days post inoculation, and quantification as % of plasma cells expressing Ki67. **(H)** Somatic hypermutation analysis: Pie charts show the number of mutations within the JH4 intron of GC B-cells and plasma cells from td-LNs of 4T1.2 tumor-bearing WT mice. **(I)** Accompanying table details base substitution patterns (A, T, C, G) in GC B-cells and plasma cells from td-LNs of 4T1.2 tumor-bearing WT mice. **(J)** Expression of the EF plasma cell signature in MT1X-high versus conventional plasma cells from an external single-cell RNA-seq dataset. **Statistical analysis:** Mann–Whitney U test used in (B); *p* < 0.05. Data in (E); n = 2 for PC^High^, n = 3 for PC^Low^. (G) are representative of two independent experiments. Data in (H) and (I); n = 3.

To test this hypothesis, we sought to build an EF-PC transcriptional signature. First, we set a stringent SHM cutoff using single-cell V(D)J data from wild-type vs Aicda⁻^/^⁻ B-cells, which lack activation-induced cytidine deaminase (AID) and therefore do not undergo SHM^32^ (**Fig. 2D-left**). Applying this threshold to a paired PC transcriptome-V(D)J dataset^33^ identified genes enriched in PCs with low-SHM, yielding a 65-gene EF-PC signature (**Fig. 2D-middle**). We then applied this signature to a new generated spatial transcriptomics dataset from human LNs from immune-cold TNBCs with high (PC^High^) and low (PC^Low^) PC burden (**Fig. 2D-right**). Using GeoMx DSP with immunofluorescence-guided segmentation, we isolated PCs only regions of interest (ROIs) in patient LNs by masking to CD138⁺ and excluding pan-cytokeratin⁺ regions. These PC segments expressed canonical markers (*JCHAIN*, *XBP1*, *PRDM1, MZB1, SDC1***; Ext. Fig. 2D**), validating cell identity. PCs from immune-cold tumors with high PC burden (PC^High^) were strongly enriched for the EF signature, with upregulation of cell-cycle, metabolic and inflammatory programs, including MYC targets and mitotic-spindle activity (**Fig. 2E,F**; **Ext. Fig. 2E**). Concordantly, >90% of td-LN PCs in the 4T1.2 model were Ki67⁺ (**Fig. 2G**). SHM analysis of the JH4 intron of 4T1.2 td-LN PCs revealed few mutations and low C→T transition frequency, compared to GC B-cells (**Fig. 2H,I**), consistent with limited AID activity and an EF origin. Together, these data indicate that the PCs in LNs of immune-cold TNBC tumors with high PC burden (PC^High^) arise predominantly via an EF pathway.

### EF-PCs link to poor prognosis across cancers

Projecting the newly defined EF-PC signature onto an orthogonal scRNA-seq pan-cancer tumor-infiltrating PC atlas revealed alignment with the MT1X-high PC cluster previously linked to poor prognosis and checkpoint-blockade resistance^19^. This was not only evident in TNBC, but also across other cancers, including Basal Cell Carcinoma, Clear Cell Renal Carcinoma, Colorectal cancer and Non-Small Cell Lung Cancer (**Fig. 2J**). This cross-cohort validation suggests that the transcriptional program we define in TNBC is not tumor-type-restricted and indicates that EF-PC differentiation is a recurrent, maladaptive immune program with prognostic and therapeutic relevance across cancers.

### EF-PCs block CD8^+^ T-cell activation in LNs

To test the functional impact of cancer-induced EF-PC hyperplasia, we depleted PCs in 4T1.2-bearing mice. Anti-CD138 given every other day for 14 days efficiently reduced EF-PCs (**Fig. 3A-D**) without altering total B-cells or GC B-cells (**Ext. Fig. 3A-D**). PC depletion curtailed tumor growth (volume and mass, **Fig. 3E**) and was accompanied by a marked increase in CD8⁺ T-cells in td-LNs (**Fig. 3F,G**). An orthogonal genetic approach to deplete EF-PCs (Jchain_CreERT2-DTA^24,25^) recapitulated these effects, reducing tumor burden with increased td-LN CD8⁺ T-cells (**Ext. Fig. 4A-D**), and pharmacologic EF-PC depletion with bortezomib similarly lowered tumor growth (**Ext. Fig. 4E-G**). Together, these interventions support a direct, suppressive role for EF-PCs on anti-tumor immunity.

**Figure 3:**
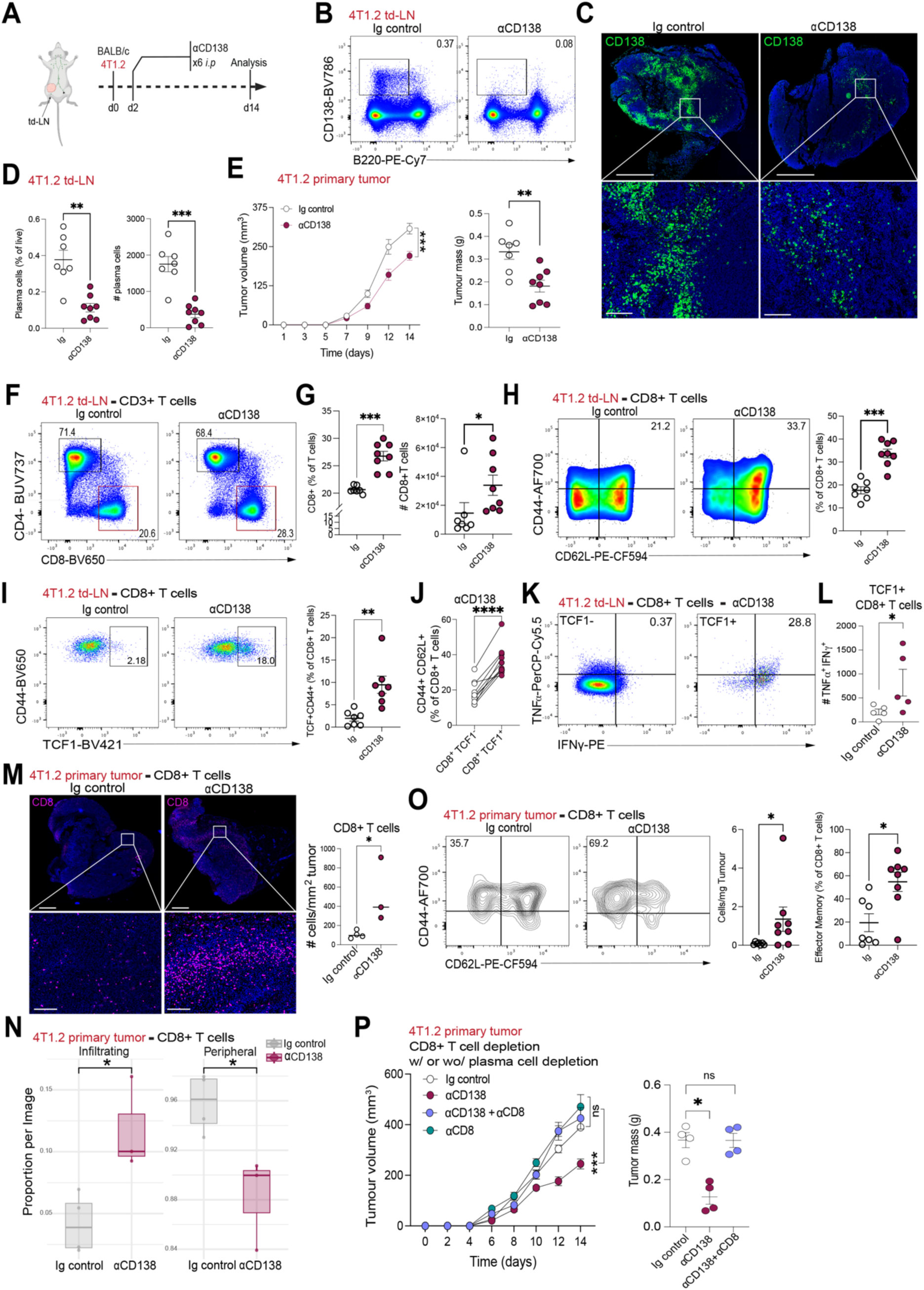
Extrafollicular-PCs inhibit stem-cell-like CD8^+^ T cell formation in LNs. **(A)** Experimental overview: BALB/c mice bearing 4T1.2 tumors were treated with an anti-CD138 monoclonal antibody (αCD138) to deplete plasma cells. **(B)** Flow cytometric analysis of plasma cells in tumor-draining lymph nodes (td-LNs) following αCD138 treatment. **(C)** Representative immunofluorescence images of CD138⁺ plasma cells in td-LNs from mice treated with Ig control or αCD138. Scale bar top panel: 500μm, scale bar bottom panel: 200μm. **(D)** Percentages and absolute numbers of plasma cells from the td-LNs of mice treated with Ig control or αCD138. **(E)** Tumor growth curves showing tumor volumes in αCD138-treated mice compared to isotype (Ig) control and tumor mass at endpoint (d14). **(F)** Flow cytometry plots of CD4⁺ and CD8⁺ T-cells in td-LNs **(G)** Quantification of % of CD8+ T-cells and absolute numbers in td-LNs of mice treated with Ig control or αCD138. **(H)** CD44 and CD62L expression on CD8⁺ T-cells in td-LNs, with quantification of CD44⁺CD62L⁺ subset. **(I)** Frequency of TCF1⁺ CD8⁺ T-cells in td-LNs as a % of CD8+ T-cells, with representative flow cytometry plots. **(J)** Co-expression of CD44 and CD62L on TCF1^−^ and TCF1⁺ CD8⁺ T-cells, shown as % of parent population. **(K)** TNF-α and IFN-γ expression in TCF1⁻ and TCF1⁺ CD8⁺ T-cells **(L)** quantification of absolute TCF1+ cytokine-producing cell numbers per td-LN. **(M)** Immunofluorescence images of CD8+ T-cell infiltration in primary tumors of mice treated with Ig control or αCD138. **(N)** Quantification of the proportion of CD8+ T-cells infiltrating the tumor bed vs restricted to the peripheral areas of the tumor **(O)** CD44 and CD62L expression on intratumoral CD8⁺ T-cells, with quantification of total CD8⁺ T-cells per mg tumor and frequency of effector memory (CD44⁺CD62L^neg^) subset. **(P)** Tumor growth kinetics and endpoint tumor mass in mice treated with Ig control, αCD138, αCD8, or the combination (αCD138 + αCD8). **Statistical analysis:** Tumor growth curves (E, P): two-way ANOVA with Bonferroni’s correction. Group comparisons (D, E, G, H, I, L, M): Mann–Whitney U test. Paired comparisons (J): Wilcoxon signed-rank test. Endpoint tumor mass (M): one-way ANOVA. *p* < 0.05 (\**), p < 0.01 (*\*\**), p < 0.0001 (***\*\*\***). Panels (D-O) are data from two independent experiments. (P) represents one experiment with 4 mice in each group.

**Figure 4:**
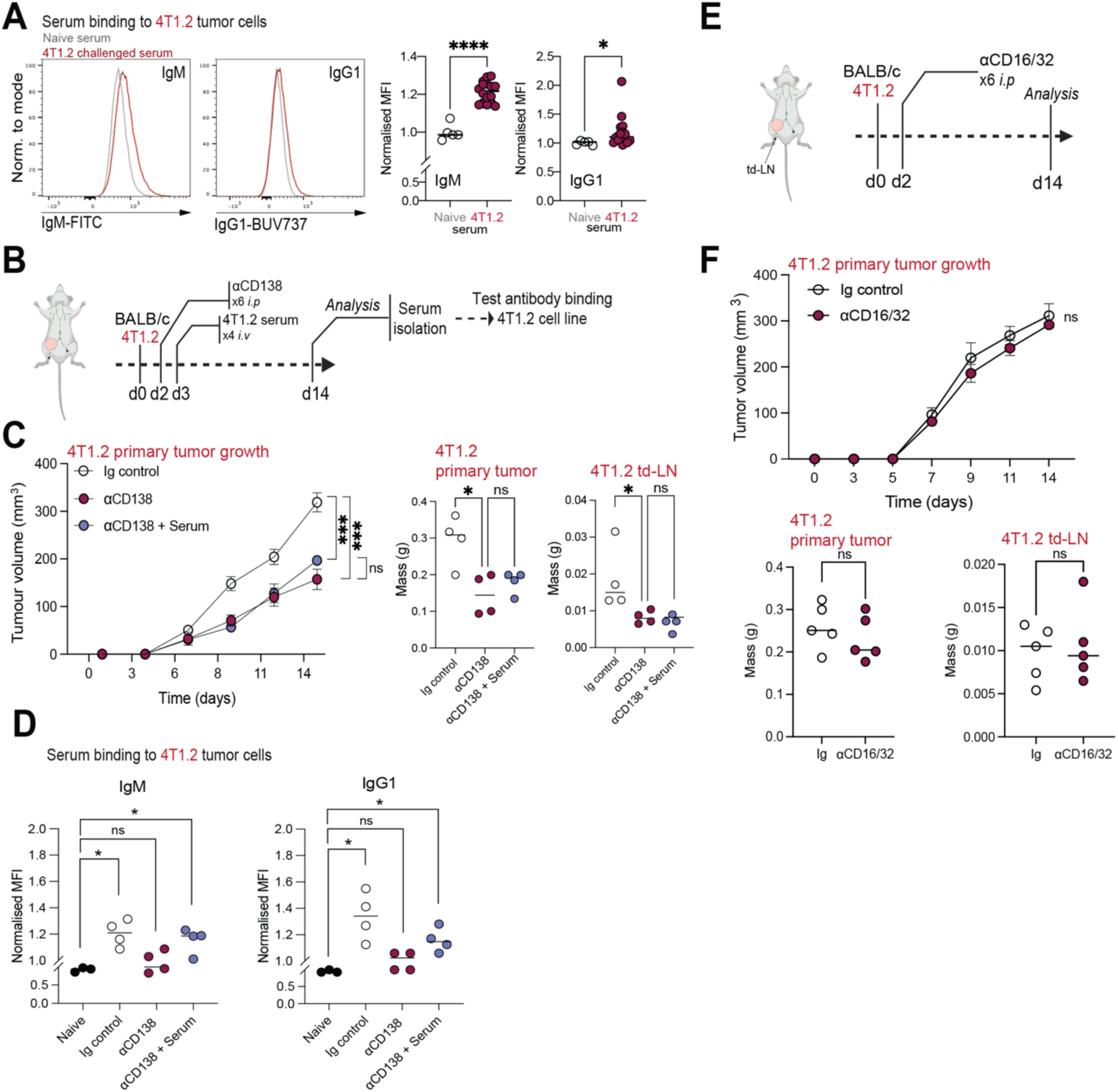
Extrafollicular-PC-mediated immunosuppression is antibody independent. **(A)** Normalized median fluorescence intensity (MFI) of IgM and IgG1 binding of naïve sera or sera from 4T1.2 tumor bearing mice to the 4T1.2 cell line, with representative flow cytometry plots **(B)** Experimental design for αCD138 antibody treatment and serum transfer experiment and subsequent isolation for antibody binding **(C)** Tumor growth curves of mice treated with Ig control, αCD138, and αCD138-treated mice that received serum from 4T1.2 tumor bearing mice, with tumor mass and td-LN mass at endpoint **(D)** Normalized median fluorescence intensity (MFI) of IgM and IgG1 binding of naïve sera or sera taken from serum transfer experiment to 4T1.2 cell line **(E)** Schematic of αCD16/32 treatment in 4T1.2 mice **(F)** Tumor growth curves of mice treated with Ig control or αCD16/32 antibody, with tumor mass and td-LN mass at endpoint **Statistical analysis:** Mann–Whitney U test was used in (C), (D), (E), (F), (G), and (K). *p* < 0.05 (*). (A) is representative data from four independent experiments. (B-F) represent one experiment with 4 mice in each group.

Phenotypic profiling revealed that EF-PC depletion resets CD8⁺ T-cell fate. In td-LNs, CD8⁺ T-cells shifted toward a central-memory phenotype (CD44⁺CD62L⁺; **Fig. 3H**) and upregulated TCF1 (**Fig. 3I,J**), defining a stem-like memory pool, which sustain long-term anti-tumor immunity^34,35^. These TCF1⁺ cells co-expressed PD-1 and CXCR5 (**Ext. Fig. 5A**) and produced more TNFα and IFNγ (**Fig. 3K,L**), consistent with a progenitor-like, tumor-reactive state. EF-PC depletion increased overall leukocyte infiltration at the tumor bed (CD45⁺; **Ext. Fig. 5B,C**), reflected by increased CD4⁺ and CD8⁺ T-cell numbers (**Fig. 3M-O**; **Ext. Fig. 5C**), and deeper parenchymal infiltration of CD8⁺ T-cells rather than peripheral accumulation (**Fig. 3M,N**). Intratumoral CD8⁺ T-cells adopted an effector-memory phenotype (CD44⁺CD62L^neg^; **Fig. 3O**).

**Figure 5:**
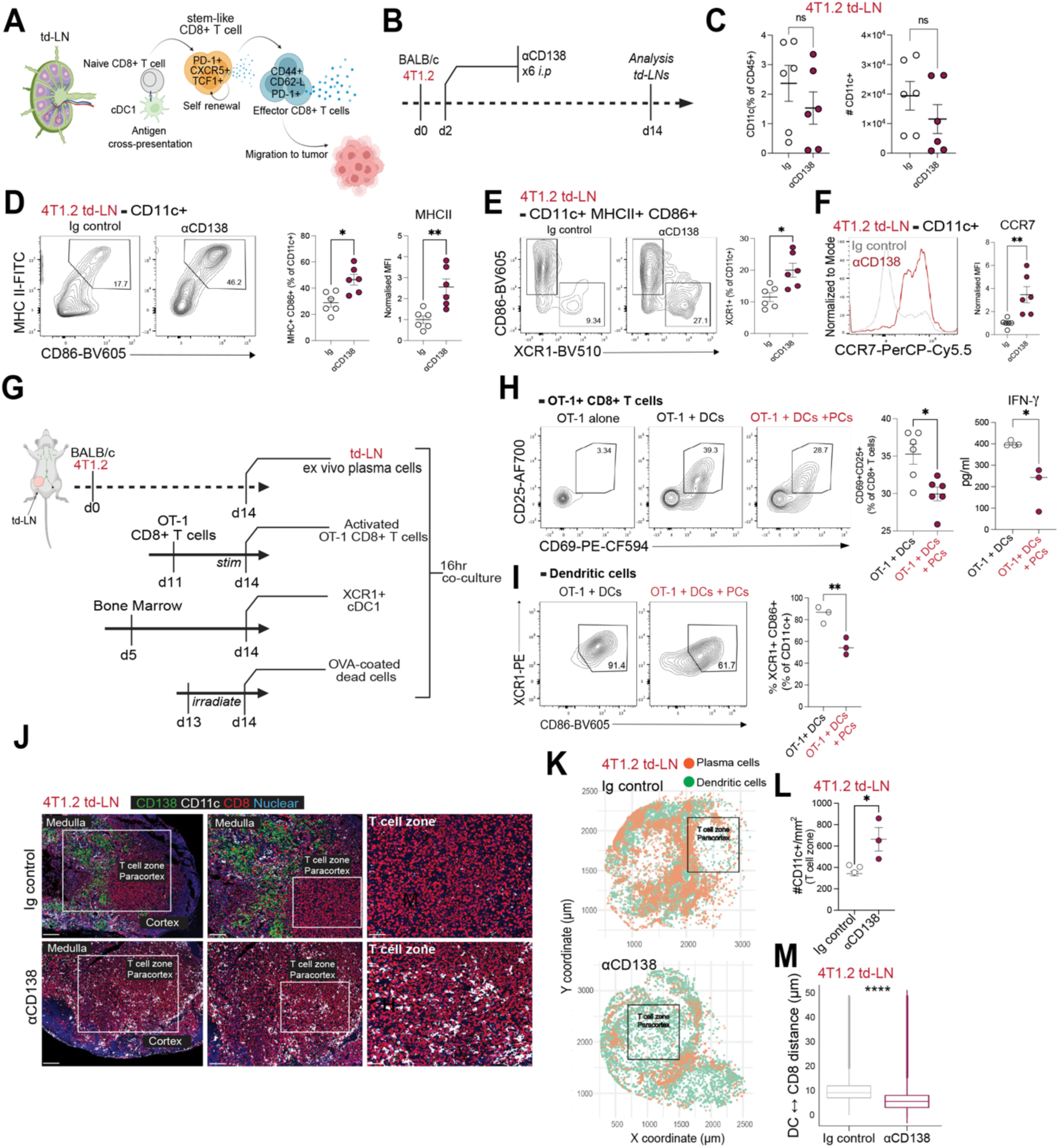
Extrafollicular-PCs hinder dendritic cell migration and activation in LNs. **(A)** Schematic illustrating the role of TCF1⁺ CD8⁺ T-cells in tumor-draining lymph nodes (td-LNs) as a reservoir for tumor-reactive effectors and the importance of antigen presentation by dendritic cells (DCs) in initiating these responses. **(B)** Experimental design for αCD138 antibody treatment in the 4T1.2 TNBC model. **(C)** Frequency and absolute number of CD11c⁺ cells (DCs) among CD45⁺ cells in td-LNs of Ig control vs αCD138-treated mice. **(D)** Expression of MHC II and CD86 on CD11c⁺ cells in td-LNs, shown as percentage of MHC II⁺CD86⁺ cells among CD11c⁺ DCs and MHC II median fluorescence intensity (MFI). **(E)** Frequency of XCR1⁺ cells among MHC II⁺CD86⁺CD11c⁺ DCs in td-LNs of Ig control vs αCD138-treated mice. **(F)** CCR7 expression (MFI) on CD11c⁺ DCs within td-LNs. **(G)** Schematic of in vitro cross-presentation assay: OT-I⁺ CD8⁺ T-cells were co-cultured with XCR1⁺ bone marrow-derived DCs and 4T1.2 tumor-associated plasma cells. **(H)** Activation of OT-I⁺ CD8⁺ T-cells with or without co-culture with plasma cells, shown by CD25 and CD69 co-expression and CD69⁺CD44⁺ T-cell frequency. IFN-γ levels in supernatants measured by ELISA. **(I)** Flow cytometry plots of XCR1+CD86+ DCs (gated on CD11c+) in wells containing OT-1 T-cells + DCs or wells containing OT-1 T-cells + DCs + PCs, with quantification of XCR1+CD86+ DCs. **(J)** Representative immunofluorescence images showing spatial localization of CD138⁺ plasma cells (green), CD11c⁺ DCs (white), and CD8⁺ T-cells (red) in td-LNs from Ig control or αCD138-treated mice. (Scale bars in left panel: 200mm, scale bars in middle panel: 100mm, scale bars in right panel: 50mm. **(K)** Spatial organization of CD138⁺ PCs and CD11c⁺ DCs within td-LNs. **(L)** Quantification of CD11c⁺ DCs per mm² within T-cell zones of td-LNs. **(M)** Distance (µm) between CD8⁺ T-cells and CD11c⁺ DCs in td-LNs, measured by image analysis. **Statistical analysis:** Mann–Whitney U test was used in (C-F), (H, I) and (K). *p* < 0.05 (*). (C-M) are data from two independent experiments.

CD8⁺ T-cells were required for therapeutic benefit. While CD8⁺ T-cell depletion alone had little effect in the immune-cold ICB-refractory 4T1.2 model^36^, it completely abrogated the tumor control achieved by EF-PC depletion (**Fig. 3P**). Thus, EF-PCs restrain the priming and expansion of TCF1⁺ stem-like CD8⁺ T-cells in td-LNs. Removing this early brake unmasks latent T-cell potential, offering a strategy to convert immune-cold tumors with high PC burden into an immune-responsive state.

### Antibody independent EF-PC-driven immunosuppression

As PCs are antibody producers, we first tested whether their tumor-promoting activity relied on antibodies. In 4T1.2-bearing mice, we detected circulating antibodies that bound native cancer cell-surface antigens, dominated by IgM with minimal IgG1 binding (**Fig. 4A**). To ask if the loss of these antibodies explains the benefit of EF-PC depletion, we transferred serum from tumor-bearing donors into EF-PC-depleted recipients. Serum transfer failed to restore tumor volume or mass (**Fig. 4B,C**), even though flow-cytometric binding assays confirmed that IgM and IgG1 binding to tumor cells was reconstituted in recipient mice (**Fig. 4D**). To exclude Fc-dependent effector functions more directly, we blocked FcγR in 4T1.2-bearing mice; however, FcγR blockade had no effect on tumor growth or mass (**Fig. 4E,F**). Together, the inability of serum transfer to rescue tumor growth, despite restored antibody binding, and the lack of efficacy of FcγR blockade argue that classical antibody mechanisms (opsonization/ADCC/ADCP) are not responsible for EF-PC-driven immunosuppression. These data point to a non-canonical, antibody-independent, cellular mode of immune control by EF-PCs.

### EF-PCs dampen cDC1 activation and CCR7-dependent migration

CD8⁺ T-cell priming within td-LNs hinges on cross-presentation by XCR1⁺ conventional type 1 dendritic cells (cDC1s), which acquire tumor antigens and license naïve CD8⁺ T-cells to enter stem-like programs before effector differentiation^37–39^ (**Fig. 5A**). We therefore asked if EF-PCs blunt this cDC1→T-cell axis.

Flow cytometry of td-LNs revealed that EF-PC depletion left total CD11c⁺ DC numbers unchanged (**Fig. 5B,C**) but markedly increased DC activation^40^, evidenced by higher fractions of CD86⁺ and MHCII⁺ cells, and increased MHCII surface expression (**Fig. 5D**). Concomitantly, the XCR1⁺ cDC1 subset was enriched (**Fig. 5E**). We also observed increased CCR7 expression on DCs upon EF-PC depletion (**Fig. 5F**), a chemokine receptor required for DC trafficking from medullary, peripheral LN regions into the T-cell zone^41^. Thus, removing EF-PCs enhances both the activation state and positional licensing of DCs that are competent for cross-presentation.

To test for direct suppression by EF-PCs, we built a cross-presentation co-culture^42^: bone-marrow-derived XCR1⁺ cDC1s were incubated overnight with pre-activated OT-I CD8⁺ T-cells^43^ and UV-irradiated 4T1.2-OVA cells, in the presence or absence ex-vivo of EF-PCs purified from td-LNs (**Fig. 5G**). EF-PCs remained present throughout the culture (**Ext. Fig. 6A**). As expected, cDC1s cross-presented OVA and robustly activated OT-I CD8⁺ T-cells (CD25, CD69, CD44 upregulation and IFNγ production; **Fig. 5H**; **Ext. Fig. 6B**). In contrast, addition of EF-PCs to the culture significantly blunted OT-I CD8⁺ T-cell activation, with reduced CD69⁺CD25⁺ and CD69⁺CD44⁺ frequencies and diminished IFNγ production (**Fig. 5H**; **Ext. Fig. 6B**). Notably, EF-PCs impaired cDC1 activation, as evidenced by reduced surface CD86 expression (**Fig. 6I**). These results demonstrate that EF-PCs directly dampen cDC1 activation and, consequently, their efficiency in CD8⁺ T-cell priming.

Spatial imaging reinforced this mechanism. In td-LNs, most CD11c⁺ DCs were found retained within medullary cords, frequently juxtaposed to EF-PCs (**Fig. 5J**). Following EF-PC depletion, DCs redistributed into the T-cell zone, which became DC-enriched and showed reduced DC-CD8⁺ T-cell separation distances (**Fig. 5K-M**). This re-patterning is consistent with CCR7-dependent migration and restores the anatomical context required for effective cDC1→CD8⁺ T-cell engagement.

## Discussion

Immune-cold (low-sTILs) TNBCs remain a major clinical problem^12,14,44^; however, their biology reflects active immunosuppression rather than immunologic inertia. We identified EF-PCs as central enforcers of this state and located the principal failure of anti-tumor priming in td-LNs.

Clinically, PC burden stratified risk within immune-cold disease. Concordant EF-PC accumulation at the tumor bed and in matched LNs specifically in patients who develop metastases indicates a coordinated, system-level program of immune suppression. This immune context dependence reconciles conflicting prognostic associations across cancers; when overall TIL infiltration is low, PCs track with cancer progression.

Mechanistically, EF-PCs function as antibody-independent suppressors. Although sera contained tumor-binding Ig, serum transfer and FcγR blockade did not reverse the anti-tumor effect of EF-PC depletion, excluding classical antibody effector pathways. Instead, EF-PCs disabled the DC1→CD8⁺ T-cell axis in td-LNs by blunting cDC1 activation and impairing CCR7-dependent repositioning to T-cell zones, thereby preventing the formation of TCF1⁺ stem-like memory CD8⁺ T-cell progenitors that underpin durable tumor control. Removing EF-PCs by antibody, genetic ablation, or pharmacological treatment, while sparing B-cell populations, reprogramed the td-LN, expanded the TCF1^+^ pool, increased intratumoral CD8^+^ T-cell effectors, and restrained tumor growth; these benefits were lost with CD8 depletion, demonstrating dependence on cytotoxic T-cell immunity. Notably, EF-PC targeting was effective in ICB-refractory tumors, placing this pathway upstream of PD-(L)1/CTLA-4 therapies and identifying EF-PCs as a tractable immunoregulatory checkpoint.

Origin and geography are central to this mechanism. EF-PCs localized to medullary cords, showed low somatic hypermutation, and exhibit a MYC-high, proliferative phenotype, features of EF differentiation and reminiscent of maladaptive responses in chronic infection and autoimmunity^16,45–48^. A low-SHM-derived EF-PC gene signature identified these cells in human td-LNs and is enriched in an MT1X⁺ PC state linked to poor prognosis and ICB-resistance across tumor types^19^, suggesting a pan-cancer immune-evasion module.

Together, these findings deliver three conceptual advances. First, they reframe PCs from passive antibody factories to active cellular gatekeepers that silence antitumor priming without antibodies. Second, the dominant checkpoint is relocated to the td-LNs, where EF-PCs were formed and enforced spatial and costimulatory separation of DCs and T-cells. Third, we define an actionable EF-PC→cDC1 checkpoint controlling access of tumors to effective T-cell immunity.

The translational path is immediate. Patient stratification could pair a simple histologic readout (CD138/BLIMP1/Ki67⁺ PCs in medullary cords) to identify immune-cold PC-high/low-sTIL tumors unlikely to benefit from ICB-alone. Therapeutic repurposing of PC–directed agents approved for Multiple Myeloma (e.g., proteasome inhibitors; anti-BCMA or anti-CD38 modalities) could debulk EF-PCs to seed a TCF1⁺ CD8^+^ T-cell pool before or alongside ICB. Mechanism-based partners that amplify cDC1 function/positioning are rational combinations.

In summary, we identified an immunosuppressive EF-PC→cDC1→T-cell cell circuit that is druggable, opening a path to reinvigorate anti-tumor immunity in poor-prognosis immune-cold TNBC and other immune-excluded cancers.

## Materials and Methods

### Patient Sample Collection

Microarray data were obtained from 124 treatment-naïve patients with invasive triple-negative breast cancer (TNBC; ER-negative, HER2-negative by IHC) treated at Guy’s Hospital, London, UK, between 1984 and 2002. For immunofluorescence analyses, primary breast tissue and involved lymph nodes (in-LNs) were collected from treatment-naïve TNBC patients through Tianjin Medical University Cancer Institute. All tissue procurement was approved by the relevant research ethics committees (KHP Cancer Biobank REC ref: 18/EE/0025; Tianjin Medical University Cancer Institute and Hospital, EK2020021). Formalin-fixed paraffin-embedded (FFPE) tissue blocks from 16 TNBC patients with low stromal tumor-infiltrating lymphocytes (sTILs ≤10%) were included. Tissue sections (5 μm) were prepared at the Tianjin Medical University Cancer Institute. TMA sections for analysis of the SCAN-B cohort (see PMID: 33568222 for TMA details) were performed at the Division of Oncology, Department of Clinical Sciences Lund, Lund University, Sweden using the Roche Ventana, and imaged using a Vectra Polaris scanner.

### Immunofluorescence Staining and Image Analysis

Tissue sections were baked at 60 °C for 1 hour, followed by dewaxing and rehydration using the Tissue-Tek DRS 2000 system (Sakura). Heat-induced antigen retrieval was then performed in citrate buffer (pH 6.0; DAKO) at 125 °C and 27 PSI for 2 minutes. After cooling to room temperature, slides were washed twice with PBS (Gibco) and air-dried. A hydrophobic barrier was drawn around each tissue section using a PAP pen (Abcam). Sections were then blocked for 30 minutes at room temperature in a humidified chamber using PBS supplemented with 5% donkey serum (Jackson ImmunoResearch) and 0.5% Triton X-100 (Sigma-Aldrich). Human primary antibodies used: CD138 (clone B-A38; Novus Biologics), CD27 (clone EPR8569; Abcam), CD20(polyclonal, Abcam), BLIMP1(EPR16655, Abcam), Pan Cytokeratin (clone AE-1/AE-3, Novus Biologics). Primary antibodies were diluted in blocking buffer and applied to tissue sections overnight at 4 °C. Mouse primary antibodies used were: CD138 (polyclonal, R&D Systems), CD11c (clone D1V9Y; Cell Signaling Technology), CD8α (clone 4SM15; ThermoFisher Scientific).

Secondary antibodies used: Donkey anti-mouse IgG-Alexa Fluor 647 (polyclonal; Jackson ImmunoResearch), Donkey anti-rabbit IgG-Alexa Fluor 488 (polyclonal; Jackson ImmunoResearch), Donkey anti-goat IgG-Alexa Fluor 594 (polyclonal; Jackson ImmunoResearch). The following day, slides were washed and incubated with secondary antibody mixes for 2 hours at room temperature (RT) in the dark. Nuclei were counterstained with DAPI (1:2000; Cell Signaling) for 5 minutes, followed by three washes in PBS. Slides were mounted using Millipore mounting medium. Whole-slide imaging was performed using the Olympus VS120 system, capturing nuclear staining at 4× magnification and multi-channel fluorescence images at 20× magnification. Image analysis was conducted using Olympus OlyVIA, QuPath^49^, exporting measurements for distance and proximity analysis in Rstudio v. 2025.05.1+513.

### Microarray and RNAseq Analyses

To define patients with “high” and “low” sTILs infiltration, a median cutoff were applied to the semi-histological scoring of sTILs in the Guy’s and SCANB cohort, at >=3 and >=20 respectively. For the METABRIC cohort, immune infiltration scores were computed using the ESTIMATE algorithm (v1.0.13), as described by Lehmann et al. (2021) *<u>Nat. Commun</u>*. Thresholds at -380 and 1634 were determined by the peak of the unimodal distribution for the METABRIC cohort respectively. Signatures for B-cell subsets were derived from public scRNAseq data from human tonsil B-cells^26^. Single sample Gene Set Enrichment Analysis (ssGSEA) module from GenePattern^50^ was employed to calculate the enrichment score of naïve, memory, light zone, dark zone B and plasma cells from normalized gene expression for individual cohorts. To classify patients into “high” and “low” groups for each B-cell subset, an optimal threshold within each enrichment score’s interquartile range was determined by the minimum p-value from the log-likelihood ratio test (LRT) in each multivariate cox-regression model, adjusted solely for sTILs group. Separate multivariate cox-regressions were repeated for each B-cell subset group fitted with other clinical covariates including age group (≥50), tumor size, tumor grade, tumor stage, nodal stage and treatment status in each cohort, with and without separation by sTILs group.

### Mouse TNBC Tumor Cells

The 4T1.2 cell line was maintained in RPMI 1640 (Gibco), supplemented with 10% fetal bovine serum (FBS) and 1% penicillin–streptomycin. Cells were passaged twice weekly upon reaching approximately 70% confluency.

### Mice

C57BL/6 *Jchain*_creERT2 animals were crossed with BALB/c mice to generate F1 offspring. BALB/c wild-type or *Jchain*_creERT2 (aged 7–12 weeks) were anesthetized with isoflurane, and 4T1.2 tumor cells were injected into the mammary fat pad in a 1:1 mixture with Matrigel. Tumor growth was monitored, and mice were sacrificed at designated time points for downstream analysis. Animal experiments were carried out in accordance with national and institutional guidelines for animal care and were approved by the Francis Crick Institute biological resources facility strategic oversight committee (incorporating the Animal Welfare and Ethical Review Body) and by the Home Office, UK.

### *In Vivo* Antibody Depletion

To investigate the roles of CD138⁺ plasma cells and CD8⁺ T-cells in the 4T1.2 tumor model, *in vivo* depletion studies were performed using monoclonal antibodies. Mice received intraperitoneal injections of anti-CD138 (clone 281-2; BioLegend) or Rat IgG2a κ isotype control (clone RTK2758; BioLegend), anti-CD8 (clone 2.43; BioXcell) or Rat IgG2b κ isotype control (clone LFT-2; BioXcell) antibodies according to the experimental schedule. Anti-CD138 antibodies were administered via intraperitoneal injection every two days, starting on day 2 post-tumor implantation. Anti-CD8 antibody was administered every four days, beginning on day 5 post-implantation. Control animals received isotype-matched control antibodies on matching schedules. Depletion efficiency was assessed by flow cytometry of lymph nodes and tumor-infiltrating lymphocytes.

### Organ Harvesting and Cell Staining

At days 7 and 14 post-injections, mice were euthanized, and tumors and lymph nodes were harvested. Organs were weighed and lymph nodes processed into single cell suspensions. Tumors were cut up into small pieces and dissociated using the mouse tumor dissociation kit (Miltenyi) according to manufacturer’s instructions to study tumor infiltrating lymphocytes. Tumor cells were then enriched for CD45+ cells using Mouse CD45 MicroBeads, using LS columns for negative selection.

The following anti-mouse antibodies were used for flow cytometric analysis: CD45(clone 104; BioLegend), CD19(clone 6D5; BioLegend), B220(clone RA3-6B2; BioLegend), CD138(polyclonal, R&D systems),CD38(clone 90; BioLegend), CD95/Fas(clone Jo2; BD Biosciences),IgM(clone II/41; BD Biosciences),IgG1(clone A85-1; BD Biosciences),CD3(clone 145-2C11; BioLegend), CD4(clone RM4-5, BD Biosciences),CD8(clone 53-6.7;BD Biosciences), CD44(clone IM7; BioLegend), CD62L(clone MEL-4; BD Biosciences), TCF1(clone S33-966; BD Biosciences),TNF(MP6-XT22;Invitrogen), IFN-γ (XMG1.2;eBioscence), CD11c(clone N418;BioLegend), MHC II (clone M5/114.15.2, Invitrogen) CD86(clone GL-1, BioLegend), CD11b(clone M1/70; BioLegend), XCR1(clone ZET; BioLegend), CD69(clone H1.2F3; eBioscience),CD25(clone PC61.5; ThermoFisher Scientific), CCR7(clone 4B12; BioLegend), NKp46(clone 29A1.4; R&D Systems).

For all staining, cells were incubated for 15 mins at 4°C in PBS with Zombie NIR live dead stain (BioLegend), washed and incubated for 20 mins with anti-mouse CD16/32 (Fc block) at 4 °C in FACs buffer (PBS + 2% + 2mM EDTA). Cells were then washed an incubated with antibodies for surface staining for 30 mins at 4 °C in FACs buffer. To study intracellular cytokine production, cells were stimulated for 4 hours at 37 °C with a Cell Stimulation Cocktail Plus (eBioscience). For intracellular staining of transcription factors, after surface antibody staining, the cells were fixed and permeabilized using the FoxP3/Transcription Factor Staining Buffer Kit (eBioscience) according to the manufacturer’s instructions. For intracellular staining of cytokine production, cells were fixed and permeabilized using the CytoFix/CytoPerm Kit (BD) according to the manufacturer’s instructions.

### Cross Presentation Assay

Mice were inoculated with the 4T1.2 cell line and tumors allowed to grow for 14 days. During this time, bone marrow was harvested from one healthy C57BL/6 mouse and cultured in RPMI 1640 supplemented with 10% FCS, 1mM sodium pyruvate, 0.1mM NEAA, 50μM β-mercaptoethanol and 1% penicillin/streptomycin supplemented with 150ng/ml FLT3L (R&D). Cells were left to differentiate for 9 days. OT-1 CD8+ cells were isolated from the spleens of OT-1-Rag1^−/-^mice and cultured with 100U IL-2 and 200pM SIINFEKL OVA peptide for 3 days in a T75 flask. The day before harvest, 4T1.2 cells were irradiated to the point of death using a UVC crosslinker (Amersham Biosciences) and left to rest overnight in PBS. The cells were then incubated with 10mg/ml OVA at 37°C for 2 hours and washed 5 times to remove excess OVA. On the day of harvest, lymph nodes from the 4T1.2 tumor bearing mice were harvested, crushed to single cell suspension, and antibodies for CD19, B220, CD138 and CD45 were added to the cells, before sorting for CD138+B220^low^ plasma cells using a BD FACSAria Fusion cell sorter. Bone marrow derived dendritic cells were incubated with an XCR1-biotin antibody (clone ZET) in 200μl MACS buffer for 10 minutes on ice. Cells were washed and then incubated with 40 μl anti-biotin Microbeads for 2 × 10^7^ cells in 200μl MACS buffer for 15 minutes at 4°C, washed and then XCR1+DCs isolated using a Miltenyi LS column. OT-1+CD8+ T-cells, XCR1+cDC1s and plasma cells.

### ELISA assay

Supernatant from the cross-presentation assay was harvested and stored at - 80°C until a later date. On the day of the ELISA, supernatants were thawed on ice. The ELISA for IFN-γ (cat: BMS606-2; Invitrogen) was performed according to the manufacturer’s instructions, and the supernatants were used undiluted.

### Mouse Serum Collection

At the experimental endpoint, blood was collected via cardiac puncture from mice treated with either isotype control or anti-CD138 depleting antibodies. Samples were left to clot at room temperature for 30 minutes, followed by centrifugation at 15,000 × *g* for 5 minutes at 4 °C. The resulting serum was carefully isolated, aliquoted to avoid repeated freeze–thaw cycles and stored at –80 °C until use in subsequent experiments.

### Serum Binding Experiments

For antibody binding experiments, 4T1.2 cells were incubated with sera from 4T1.2 tumor bearing mice diluted 1:50 in PBS for 30 minutes at room temperature, washed with FACs buffer, and stained with fluorescently labelled antibodies against mouse IgM and IgG1 for 30 minutes at 4°C.

### Flow Cytometry and Data Analysis

Samples were acquired on a BD LSRFortessa flow cytometer, and data were recorded in FCS 3.0 format using BD FACSDiva software. Flow cytometry data were analyzed using FlowJo (v10.7.2).

### Extrafollicular Signature Derivation

The extrafollicular gene signature was generated based on somatic hypermutation (SHM) levels on the B-cell receptor (BCR) sequences, quantified by Shazam package implemented in the Dandelion (v0.3.2) pipeline. By comparing cells between wild-type and Aicda^−/-^ mice^26^, a manual SHM threshold (mu_freq = 0.00075) was adopted to distinguish plasma cells with high and low SHM^27^. Differential expression analysis was performed, and the top 100 significantly upregulated genes (padj < 0.05) in low SHM splenic plasma cells were selected to define the signature. This signature was subsequently mapped to human datasets, and using available human genes was refined to a 65-gene signature for downstream analysis.

### Single-cell RNA-seq analysis of plasma cell states

We used the publicly available single-cell RNA sequencing (scRNA-seq) dataset from Fitzsimons et al., “A pan-cancer single-cell RNA-seq atlas of intratumoral B and plasma cells” (Cancer Cell, 2024). The integrated Seurat object (BCELL_integrated.rds) was loaded, which includes cells from multiple cancer types and annotated plasma / B-cell subsets. Only treatment naïve tumor settings were included. Seurat’s AddModuleScore was used to compute EF signature scores for each cell. Cells were then subset to plasma cell clusters (i.e. MT1X-high, conventional). For each cancer type, EF signature scores were compared between MT1X-high vs conventional plasma cell populations using Wilcoxon rank-sum tests, with Bonferroni correction for multiple comparisons. P values were corrected for multiple comparisons using the Bonferroni method.

### JH4 Intron Analysis

Quality control of the fastq files for JH4 intron reads was performed using DADA2 (v1.28.0) in R, where reads were trimmed to 240bp using filterAndTrim() function with maxEE set to c(2,2). The output forward and reverse reads were merged into a single amplicon sequence using bbmerge from bbmap (v39.01) with default settings, aligned to the JH4 reference sequence of 433bp for BALB/cJ mouse using Mafft (v7.475). Samples containing < 400 reads, as well as sequences containing insertion/deletions were excluded from subsequent analysis. The frequency of single nucleotide polymorphisms (SNPs) for each cell type/mouse model was quantified as average counts per million (CPM) across replicates. The sequence was taken from BALB_cJ_v1 reference genome using Ensembl 103.

### Spatial Transcriptomics GeoMx DSP

Formalin-fixed paraffin-embedded (FFPE) TNBC tissue sections (5 μm) were processed for spatial transcriptomic profiling using the NanoString GeoMx® Digital Spatial Profiler (DSP). Sections were deparaffinized, rehydrated, and subjected to heat-induced antigen retrieval in citrate buffer (pH 6.0; DAKO). Slides were then incubated with a panel of morphology markers, including Syto13 (nuclei), pan-cytokeratin (panCK; epithelial cells) and CD138 (plasma cells) alongside oligonucleotide-tagged RNA probes targeting the whole transcriptome. Regions of interest (ROIs) were selected based on morphology marker staining (CD138+panCK-) and captured via UV-mediated oligonucleotide release. Released oligos were collected, libraries assembled. Data processing, including quality control and normalization, was carried out using an in-house pipeline.

### Tumour immunofluorescence analyses

Cell-level spatial data were generated by cell segmentation and classification performed in QuPath (v0.6.0-x64) and exported for tumour sections (Tumour_001–Tumour_007) as tab-delimited files containing cell classifications and centroid coordinates (µm). Data were analysed in Rstudio using *dplyr* and *ggplot2*. Column names were standardised using make.names(), and cells annotated as “CD8” were classified as CD8⁺ T cells, with all others designated as non-CD8 cells. For each image, the tumour centre was defined as the mean X and Y centroid coordinates of unlabelled cells (cells lacking a class annotation), representing the tumour bulk. The Euclidean distance of each CD8⁺ T cell to the tumour centre was calculated and normalised to tumour size by dividing by the maximum distance of tumour cells from the centre, yielding a relative scale from 0 (centre) to 1 (edge). Images were assigned to experimental conditions (Ig control: Tumour_001, _003, _005, _006; CD138-depleted: Tumour_002, _004, _007). For each tumour, the mean normalised distance of CD8⁺ T cells was calculated as a measure of infiltration. CD8⁺ T cells were further classified as infiltrating (normalised distance ≤0.33) or peripheral (>0.33), and proportions were calculated per image.

### Lymph node immunofluorescence analyses

Cell segmentation and classification were performed in QuPath (v0.6.0-x64) on immunofluorescence-stained lymph node sections, and cell-level data were exported as .csv files containing centroid coordinates and assigned cell classes. Cells were classified based on CD8⁺, CD11c⁺ and CD138⁺ staining. Spatial distributions of CD11c⁺ and CD138⁺ cells were visualised using centroid coordinate maps. Data were imported into R and analysed using *tidyverse*, *dplyr* and *ggplot2*. Centroid coordinate columns were standardised after import, missing class annotations were reassigned as “Unclassified”, and rows containing ambiguous class labels were excluded. Cells classified as CD8⁺ T cells and CD11c⁺ cells were extracted from each image, and Euclidean distances from each CD8⁺ T cell to the nearest CD11c⁺ cell were calculated based on centroid coordinates. Nearest-neighbour distances were computed for each CD8⁺ T cell and summarised per image. In addition, CD8⁺ T cells were manually quantified in QuPath within T cell zone regions defined by CD8 staining, and counts were normalised to the area of the T cell zone (mm²).

### Statistical Analysis

All statistical analyses were conducted using GraphPad Prism (Version 9) and RStudio v.2025.05.1+513. A p-value < 0.05 was considered statistically significant.

## Acknowledgements

We thank Aleksey Chudnovskiy [Francis Crick Institute (FCI), London, UK], Jose M. Adrover [FCI, London, UK], the members of the Immunity and Cancer laboratory [FCI, London, UK], and of the Cancer Bioinformatics [King’s College London (KCL), London, UK] for critical discussions and comments. We thank Tony Ng [KCL, London, UK] for the 4T1.2 cell line. We thank the Cancer Biobank [KCL, London, UK] for their support, expertise and providing human samples for data analysis. We thank the FCI scientific platforms (Biological Resource Facility, Flow Cytometry, Histopathology, Light Microscopy, Advanced Sequencing) for expert advice and technical support. Schematics were created with Biorender. The authors would like to acknowledge patients and clinicians participating in the SCAN-B study, personnel at the central SCAN-B laboratory at the Division of Oncology, Lund University, the Swedish national breast cancer quality registry (NKBC), Regional Cancer South, RBC Syd, and the South Sweden Breast Cancer Group (SSBCG).

## Author contributions

Conceptualization: E.A., A.G., and D.P.C.

Methodology: E.A., V.B., M.S.H., I.W., M.L., M.J., A.X., J.Q., and C.C.

Investigation: E.A., V.B., and M.S.H.

Resources: A.G., F.L., C.G., J.S., and D.P.C.

Visualization: E.A., and M.S.H.

Funding acquisition: A.G., S.N.K., and D.P.C.

Supervision: A.G., S.N.K., and D.P.C.

Writing-original draft: E.A., A.G., and D.P.C.

Writing-review and editing: All authors.

## Extended data

**Extended Figure 1:**
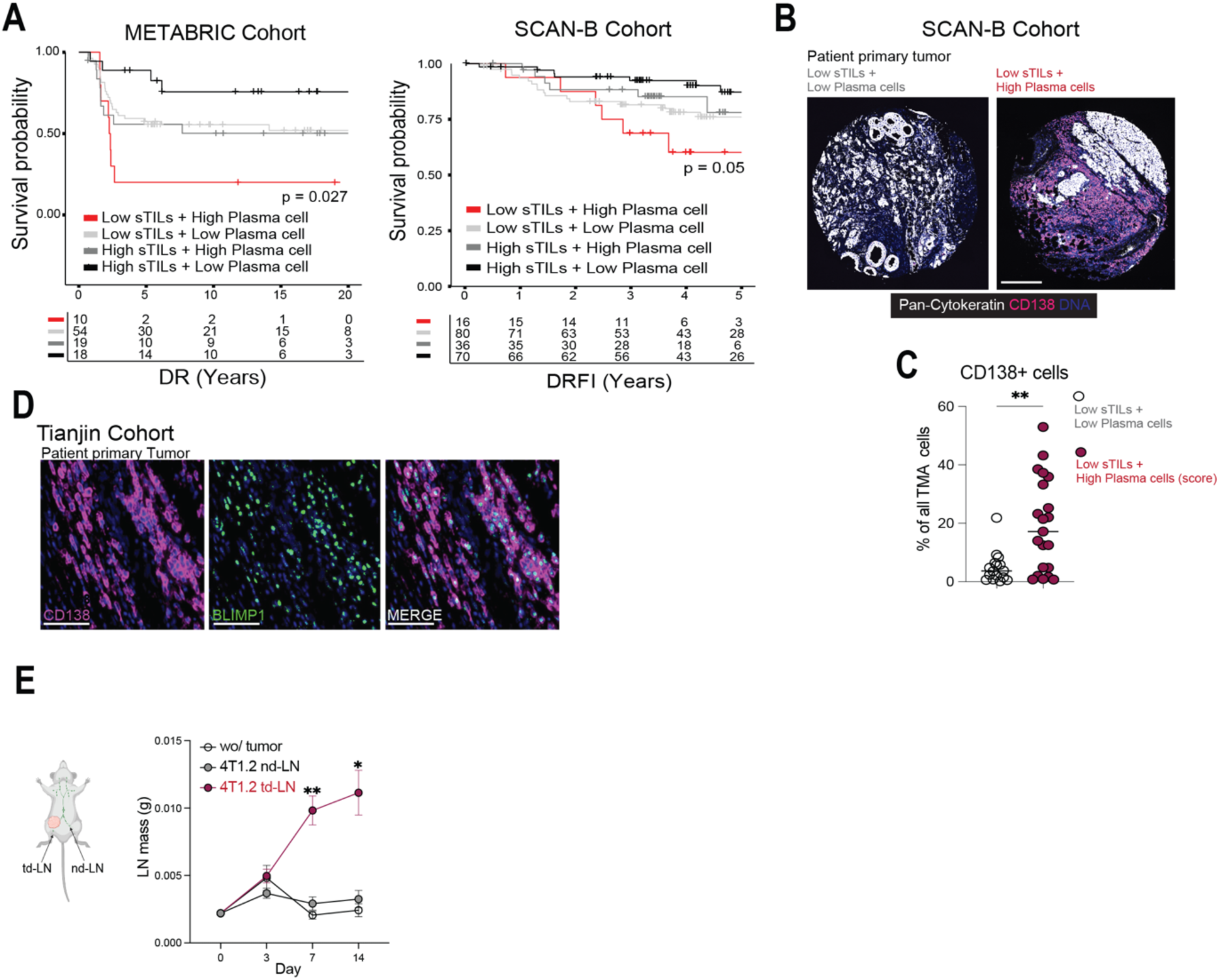
Plasma-cell burden stratifies risk in immune-cold TNBC. **(A)** Kaplan–Meier plots of distant recurrence time (DI) in the METABRIC cohort and distant recurrence-free interval (DRFI) in the SCAN-B cohort, stratified by sTIL status and plasma cell gene signature: High sTIL / high plasma cell signature (black), High sTIL / low plasma cell signature (dark grey), Low sTIL / low plasma cell signature (light grey), Low sTIL / high plasma cell signature (red). **(B)** Representative immunofluorescence images and quantification of CD138⁺ plasma cells in tumors from SCAN-B patients with either low or high plasma cell gene expression scores. Scale bar: 250mmm. **(C)** Quantitation of data analyzed in (B). **(D)** Representative staining of BLIMP1⁺CD138⁺ plasma cells in primary tumors. Scale bar: 200mm. **(E)** Mass of tumor-draining lymph nodes (td-LNs) over time in tumor-free control mice, or mice injected with 4T1.2, measured across 14 days **Statistical analysis:** Survival analyses in (A,B): Likelihood ratio test; HR = hazard ratio; CI = confidence interval. Data from (E): Mann–Whitney U test. One-way ANOVA was used in (C). *p* < 0.05 (*), *p* < 0.01 (**). Data in (E) are representative of five independent experiments.

**Extended Figure 2:**
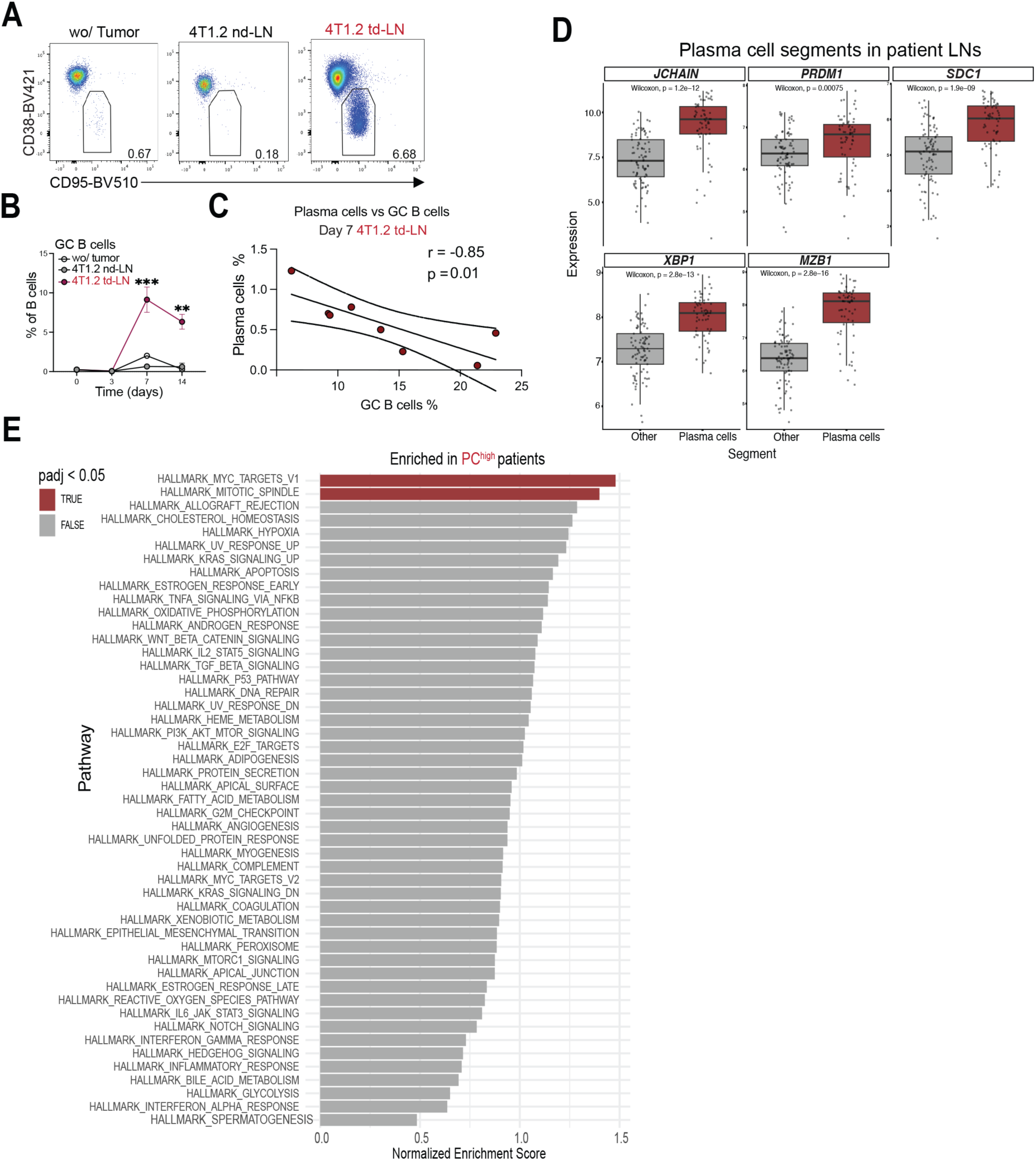
Inverse PC-GC dynamics in lymph nodes. **(A)** Representative flow cytometry plots of germinal center (GC) B-cells in td-LNs and non-draining lymph nodes (nd-LNs) from tumor-free mice or those bearing 4T1.2 tumors. **(B)** Quantification of GC B-cells (% of B-cells). **(C)** Inverse correlation between plasma cells (% of live cells) and germinal center (GC) B-cells (% of B-cells) within tumor-draining lymph nodes (td-LNs) of mice 7 days after 4T1.2 tumor inoculation. **(D)** Expression of canonical plasma cell and GC B-cell–associated genes in plasma cell-enriched regions from human lymph nodes analyzed by spatial transcriptomics. **(E)** Gene Set Enrichment Analysis (GSEA) of pathways upregulated in plasma cell segments from patients with high plasma cell infiltration (PC^high^). **Statistical analysis:** Spearman’s correlation was used for (C). Mann–Whitney U tests were used in (B). *p* < 0.05 (*), *p* < 0.01 (**). Data in (A-C) are data from two independent experiments.

**Extended Figure 3:**
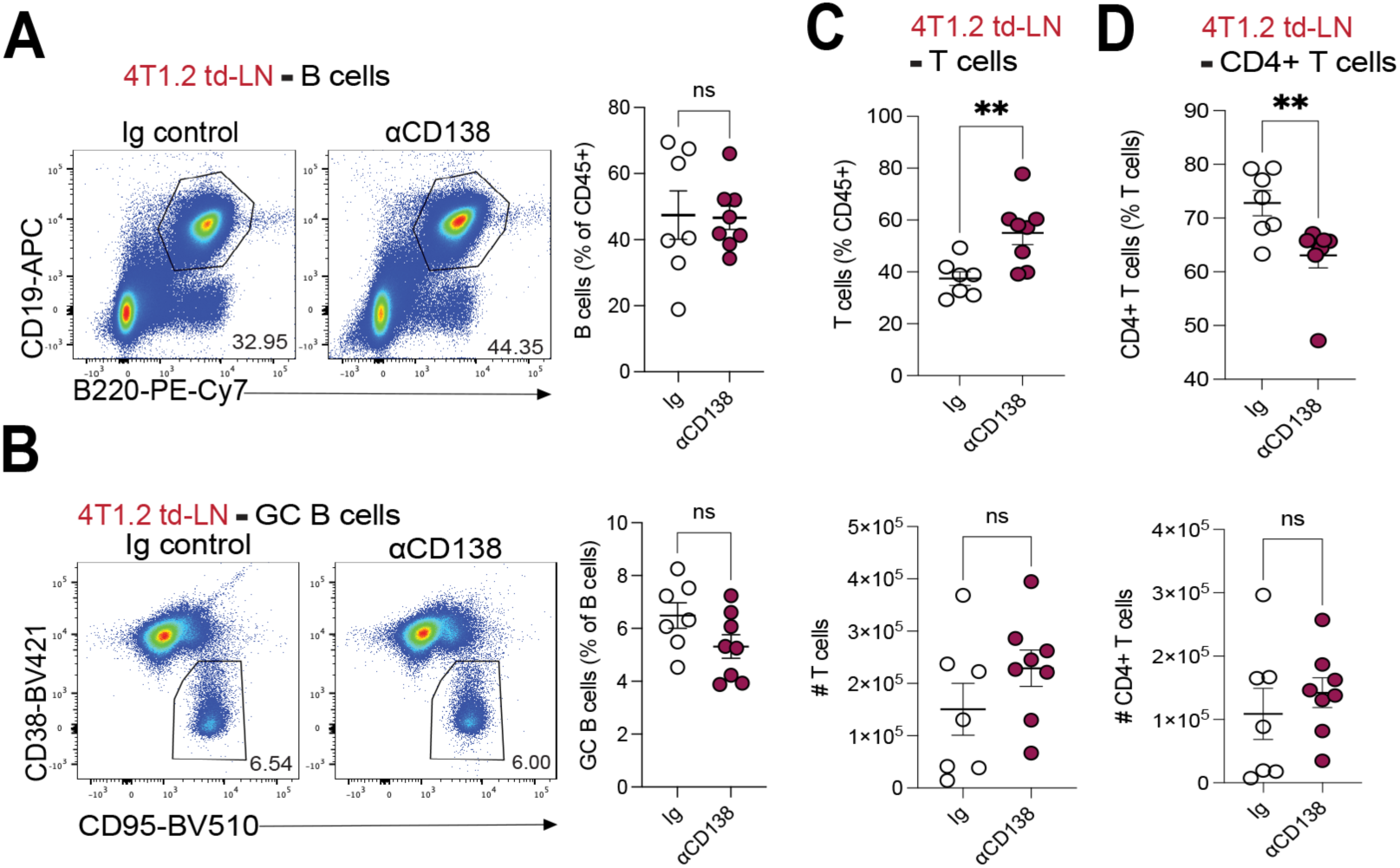
Extrafollicular-PC depletion does not impact B cell compartments in LNs. **(A)** Flow cytometric plots and quantification of the % and cell numbers of B-cells in td-LNs of mice treated with Ig control or anti-CD138. **(B)** Flow cytometric plots and quantification of the % and cell numbers of GC B-cells in td-LNs of mice treated with Ig control or anti-CD138. **(C)** Quantification of the % and cell numbers of T-cells in td-LNs of mice treated with Ig control or anti-CD138. **(D)** Quantification of the % and cell numbers of CD4^+^ T-cells in td-LNs of mice treated with Ig control or anti-CD138. **Statistical analysis:** Graphs from (A-D): Mann-Whitney U test. *p* < 0.05 (*), *p* < 0.01 (**). Data from (A-G) are data from two independent experiments.

**Extended Figure 4:**
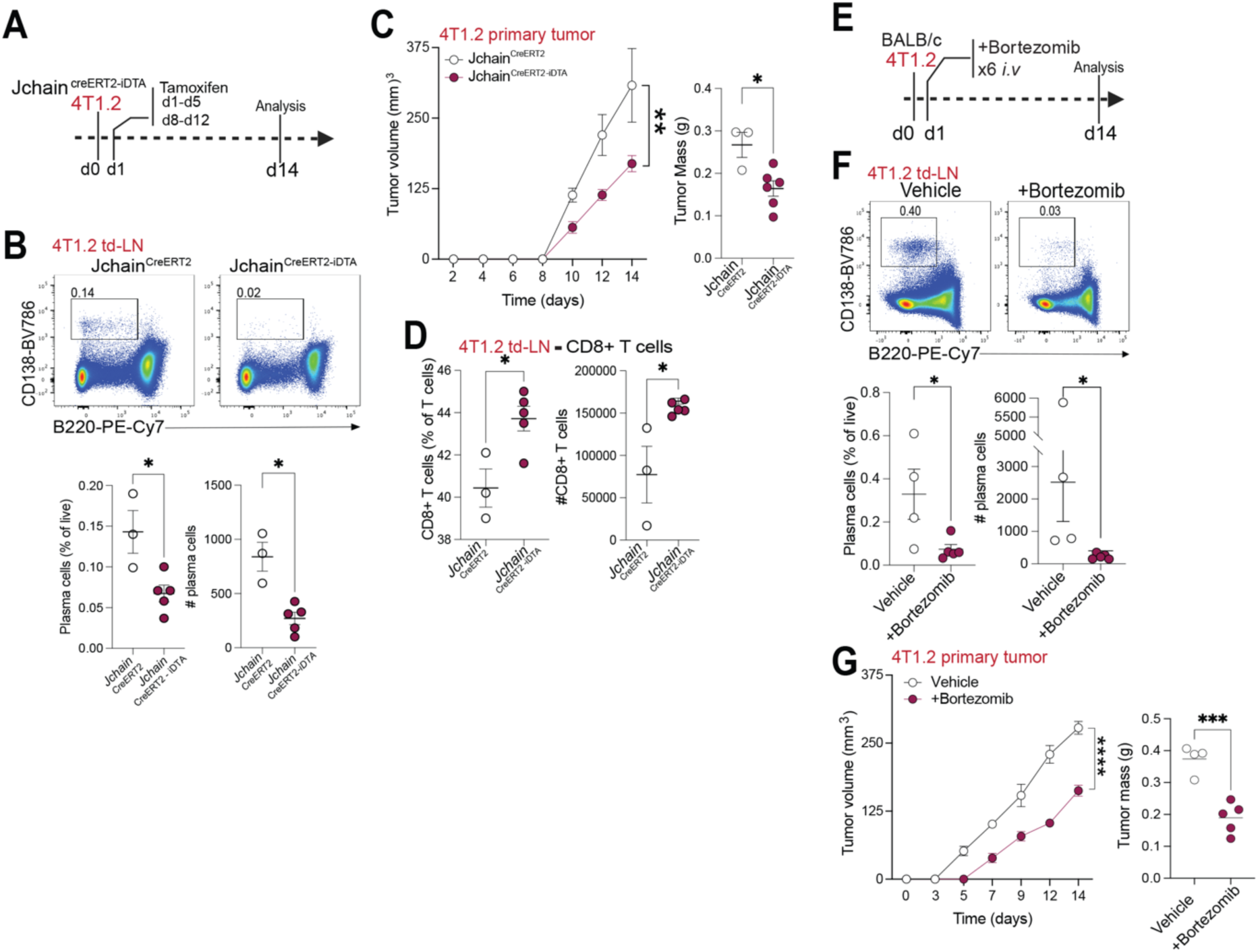
Genetic and pharmacologic depletion of extrafollicular-PCs enhances anti-tumor immunity. **(A)** Experimental schematic of tamoxifen administration in *Jchain_creERT2* and *Jchain_creERT2-DTA* mice following 4T1.2 tumor inoculation. **(B)** Representative flow cytometry plots of CD138⁺B220^neg^ plasma cells in the td-LNs of *Jchain_creERT2* and *Jchain_creERT2-DTA* mice and quantification of plasma cells in td-LNs, shown as percentage and absolute number. **(C)** Tumor growth curves and endpoint tumor mass in *Jchain_creERT2* and *Jchain_creERT2-DTA* mice. **(D)** Percentage and absolute number of CD8⁺ T-cells in td-LNs of *Jchain_creERT2* and *Jchain_creERT2-DTA* mice. **(E)** Schematic of Bortezomib treatment timeline in 4T1.2 tumor-bearing mice **(F)** Representative flow cytometry plots of CD138⁺B220^neg^ plasma cells in the td-LNs of vehicle or Bortezomib treated mice and quantification of plasma cells in td-LNs, shown as percentage and absolute number. **(G)** Tumor growth curves and endpoint tumor mass in vehicle and Bortezomib treated mice. **Statistical analysis:** Tumor growth curves in (A) and (G): two-way ANOVA with Bonferroni correction. Graphs from (B,D,F,G): Mann-Whitney U test. *p* < 0.05 (*), *p* < 0.001(***). Data from (C-I) are from one experiment with 3 to 5 mice per group.

**Extended Figure 5:**
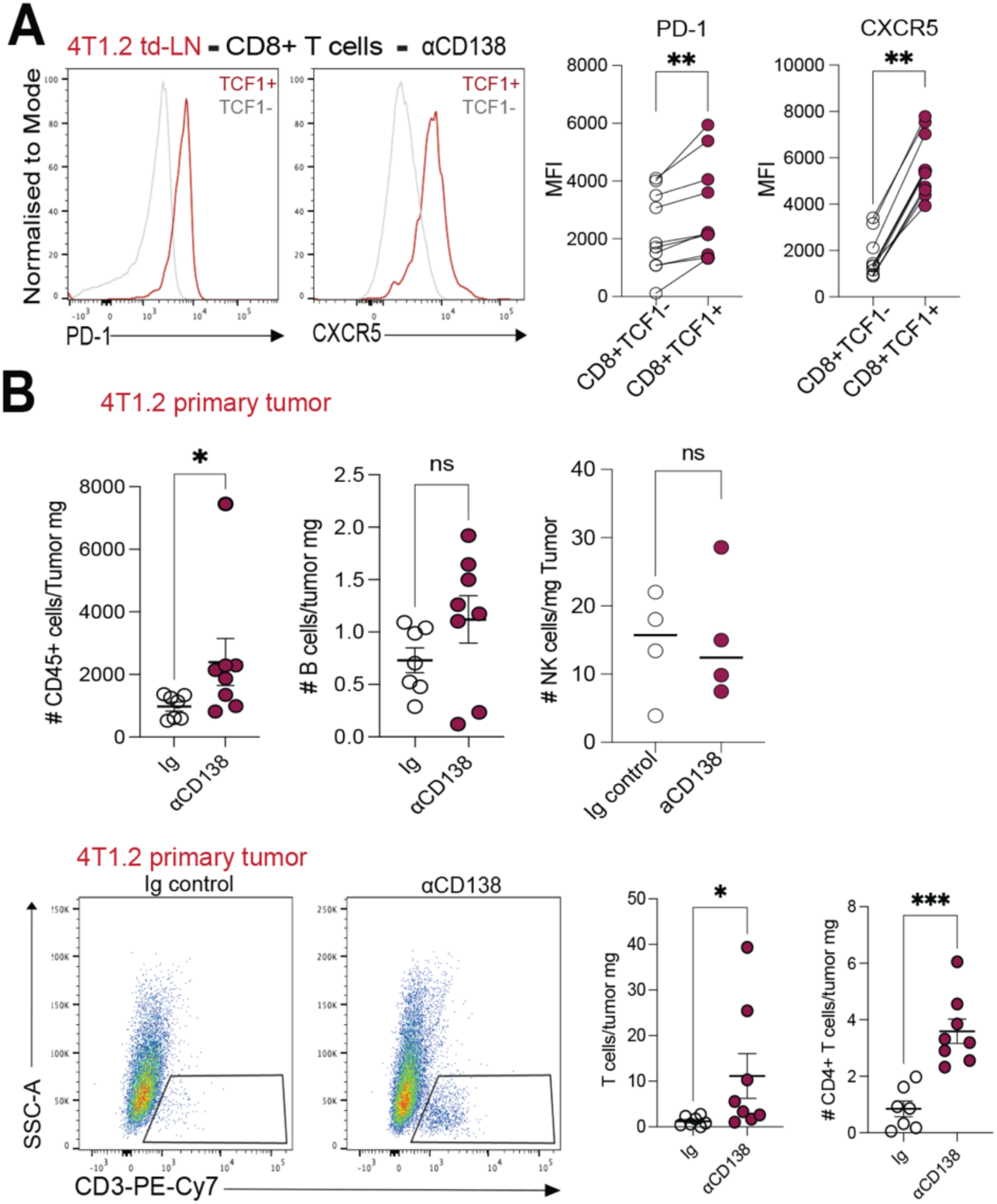
Extrafollicular-PC depletion increases intratumoral T-cell infiltration. **(A)** Flow cytometric plots of PD-1 and CXCR5 expression on TCF1- and TCF1+ CD8+ T-cells, with quantification of MFI. **(B)** CD45+ cells, B-cells and NK cells per mg of tumor in Ig control vs in αCD138- treated mice **(C)** Flow cytometry of tumor-infiltrating CD3⁺ T-cells, with quantification of total T-cells and CD4⁺ T-cells per mg of tumor in Ig control vs in αCD138- treated mice **Statistical analysis:** Graphs from (A): Paired Wilcoxon test. Graphs from (B) Mann-Whitney U test. *p* < 0.05 (*), *p* < 0.01 (**), *p* < 0.001 (***). Data from (A-B) are data from two independent experiments.

**Extended Figure 6:**
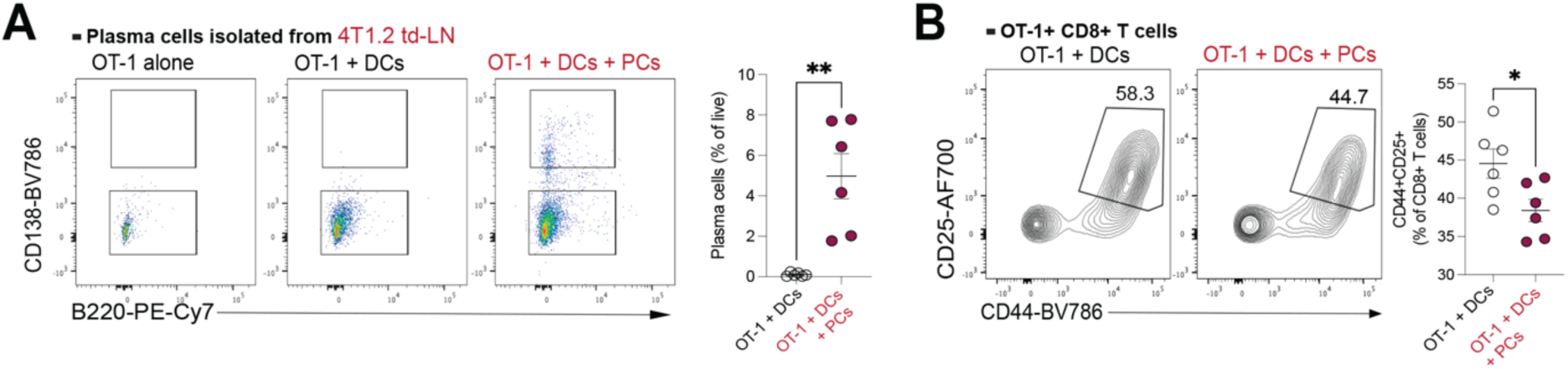
EF-PCs impair dendritic cell activation. **(A)** Flow cytometry plots of CD138+B220- plasma cells in wells containing OT-1 T-cells, OT-1 + dendritic cells (DCs) and OT-1 + DCs and plasma cells (PCs with quantification of % of plasma cells. **(B)** Flow cytometry plots of CD25+CD44+ OT-1 T-cells in wells containing OT-1 T-cells + DCs and OT-1 T-cells + DCs + PCs with quantification of % of CD25+CD44+ OT-1 cells. **Statistical analysis:** All graphs in (A-B): Mann-Whitney U test. *p <* 0.05 (*), *p* < 0.01 (**). Data are from two independent experiments.

**Extended Figure 7:**
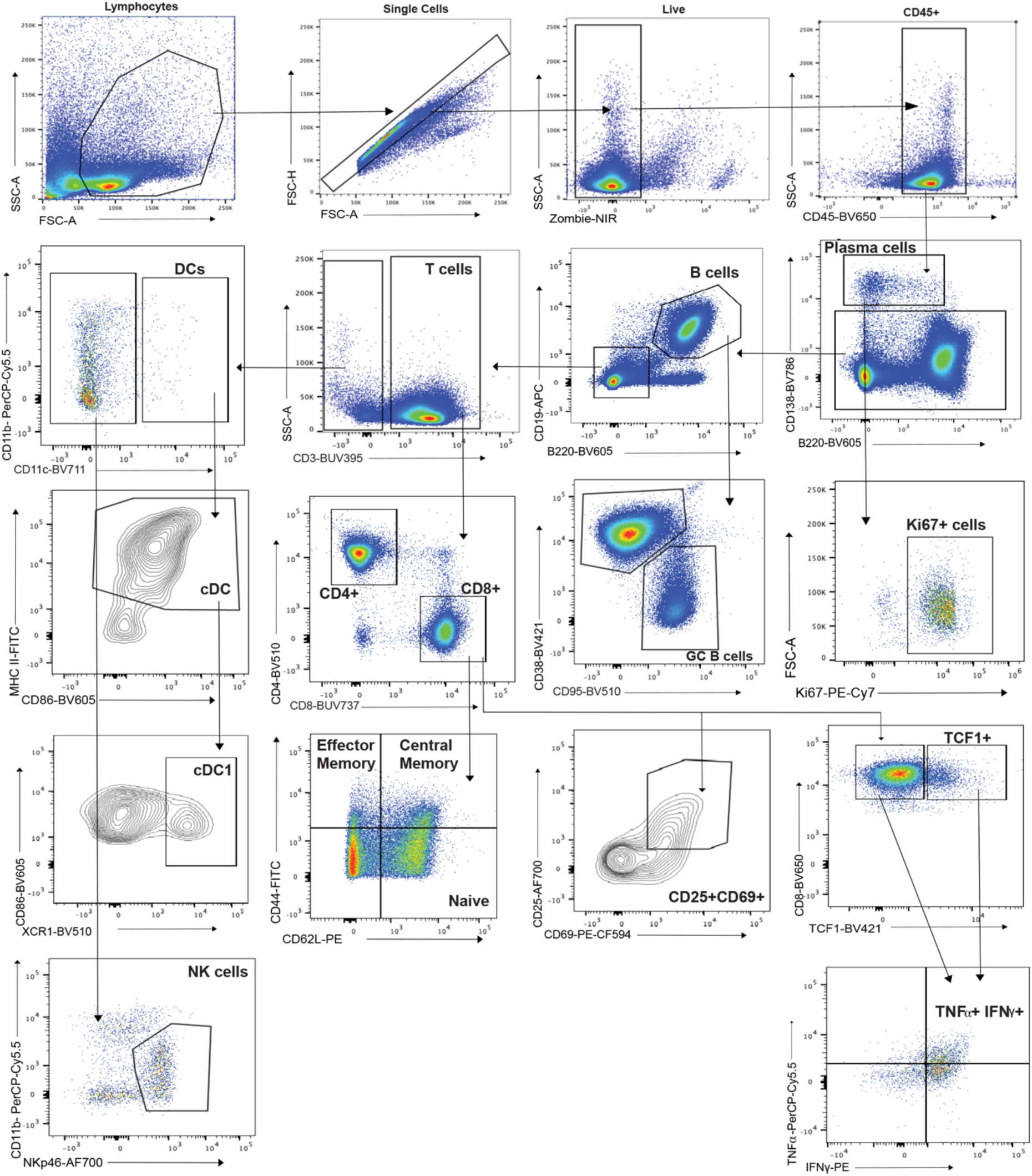
Flow cytometry gating strategy. Lymphocytes were selected by FSC/SSC, doublets excluded, and live CD45^+^ cells gated. Plasma cells (CD138^+^B220^−^) were resolved from the CD45^+^ gate with all CD19^+^B220^+^ B-cells gated on the population excluded from the plasma cell gate. Ki67+ cells were gated on plasma cells to examine proliferation. CD38^high^CD95^low^ B-cells were identified as GC B-cells. T-cells were observed from the CD19-B220- fraction as CD3+, further split into CD4+ and CD8+. CD8+ T-cells were further gated into CD44 and CD62L quadrants to identify effector memory, central memory and naïve populations. TCF1+ and TCF1- cells were gated on the parent CD8+ T-cell population and downstream on IFNg and TNFa. For *in vitro* studies, CD25 and CD69 was used to identify activation on CD8+ T-cells. DC populations were resolved from the CD3-gate as CD11c+, further gating on MHC II, CD86 and XCR1. NK cells were identified as NkP46+CD11b+/-.

